# Bromodomain proteins as new drug target for an inborn lysosomal defect

**DOI:** 10.1101/2023.12.19.572384

**Authors:** Martina Parente, Amélie Barthelemy, Claudia Tonini, Sara Caputo, Alessandra Sacchi, Stefano Leone, Marco Segatto, Frank W. Pfrieger, Valentina Pallottini

## Abstract

Inborn errors of lysosomal function often provoke disorders presenting highly variable onset, diverse visceral, neurologic and psychiatric symptoms and reduced life spans. A prime example is Niemann-Pick type C disease (NPCD). At present, therapeutic options are limited to palliative care and disease-modifying drugs, and there is a need for new treatments. Here, we explored bromodomain and extra-terminal domain (BET) proteins as a new drug target for NPCD using patient-derived skin fibroblasts. Treatment of cells with JQ1, a prototype BET protein inhibitor, enhanced the level of NPC1 protein, diminished lysosomal expansion and cholesterol accumulation, and induced extracellular release of lysosomal components in a dose- and time-dependent manner. The effect of JQ1 on protein levels was largely independent from the patient line tested, but the extent of cholesterol reduction varied in a line-dependent manner. Lastly, JQ1 enhanced and reduced cholesterol accumulation induced by inhibition of NPC1 activity and of histone deacetylases, respectively. Taken together, our results provide further evidence for epigenetic regulation of cellular NPC1 levels and cholesterol homeostasis. Pharmacologic inhibition of bromodomain proteins should be explored as candidate therapeutic approach for NPCD and as a tool to understand basic mechanisms of lysosomal function and lipid metabolism.

## Introduction

Niemann-Pick type C disease (NPCD) is a rare, autosomal-recessive, and pan-ethnic lysosomal disorder presenting progressive and ultimately fatal neurovisceral symptoms ^1,2^. Several forms of NPCD are discerned based on the onset of neurologic symptoms ^3–5^, its estimated incidence is 1:100,000 ^1^, but the adolescent/adult form may be more frequent ^6^. The primary cause of NPCD are specific variants of *NPC intracellular cholesterol transporter 1* (*NPC1*; OMIM #257220; 95% of cases) ^7,8^ or of *NPC intracellular cholesterol transporter 2* (*NPC2*; OMIM 607625; 5% of cases) ^9^. The encoded proteins NPC1 and NPC2, which are ubiquitously expressed, reside in the membrane ^10,11^ and lumen of late endosomes ^9,12,13^, respectively. Genetic manipulations ^14^, biochemical assays ^15,16^ and structure analyses ^17–22^ indicate that the two proteins export unesterified cholesterol from the endosomal-lysosomal system. Dysfunction of either protein causes intracellular accumulation of unesterified cholesterol ^23,24^ and of other lipids ^25–29^, and impairs lysosomal ^28,30–32^ and mitochondrial function ^33–37^, and autophagy ^38–40^.

Despite considerable efforts ^41,42^, therapeutic options for NPCD are limited to symptomatic treatment and to the disease-modifying drugs N-butyl-deoxynojirimycin (OGT918, Miglustat, Zavesca) ^43,44^, arimoclomol (Miplyffa) in combination with Miglustat ^45^ and N-acetyl-L-leucine (Levacetylleucine)^46^. Here, we explored bromodomain and extra-terminal (BET) proteins as new drug target for NPCD. BET proteins bind to acetylated lysines on histones and thereby regulate gene transcription ^47^. They are explored as therapeutic drug targets ^48^ for different types of cancer ^49^ and other pathologic conditions ^50,51^ including cachexia ^52^, Duchenne muscular dystrophy ^53^, Fabry disease ^54^ and retinal inflammation due to STING activation ^55^, which contributes to neurodegeneration in NPCD ^56,57^. Recently, we discovered that JQ1, a well-characterized competitive inhibitor of BET proteins ^58^, enhances the protein content of NPC1 in cultured cells ^59^. This effect has therapeutic potential for NPCD because disease severity seems to correlate with cellular levels of NPC1 protein ^60–62^ and because some pathogenic variants of NPC1 are functional, but degraded due to misfolding ^63–68^.

Using patient-derived skin fibroblasts as preclinical cell culture model ^69^, we show that inhibition of BET proteins by JQ1 increases the levels of NPC1 and reduces pathologic changes, at least in part through lysosomal release, in a dose- and time-dependent manner. Notably, the effects of JQ1 required NPC1 activity and varied in fibroblasts from distinct patients. Together, our results support further exploration of BET protein inhibition as new therapeutic approach for NPCD and as tool to study cellular cholesterol homeostasis.

## Materials and Methods

### Cell culture and drug treatment

Human dermal fibroblasts used in this study were obtained from NIGMS Human Genetic Cell Repository (Coriell Institute for Medical Research, Camden, NJ, USA). Most experiments were performed using fibroblasts from a NPCD patient homozygous for a frequent pathogenic allele [GM18453, male, p.Ile1061Thr p.Ile1061Thr ^70^] and from a sex- and age-matched healthy donor (GM05659: male, 14 months old). Selected experiments were performed using fibroblasts from heterozygous patients carrying different allele combinations [GM00110: male, 9 years, p.Pro237Ser p.Phe740_Ser741del ^60^; GM03123: female, 9 years, p.Pro237Ser p.Ile1061Thr ^60,71^; GM17920: female, p.Pro401Thr p.Ile1061Thr ^72^; GM17921: male, 5 years, p.Pro433Leu p.Ile1061Thr ^73^]. Cells were cultured in Dulbecco’s modified Eagle medium (DMEM) containing high glucose supplemented with 5% fetal bovine serum, 1% L-glutamine, 1% sodium pyruvate, 1% non-essential amino acids, and 1% penicillin-streptomycin (all Sigma-Aldrich/Merck) at 37°C and 5% CO_2_. All experiments were performed at 60–70% cell confluency and maximally 20 passages. The drugs (+)-JQ1 (#SML1524, Sigma-Aldrich/Merck), 3-beta-[2-(diethylamine)ethoxy]androst-5-en-17-one [U18666A (U18); #S9669 Selleckchem) and suberoylanilide hydroxamic acid (SAHA, Vorinostat; #SML0061, Sigma-Aldrich/Merck) were added to primary cultures at indicated concentrations after dilution from respective stock solutions (JQ1: 3 mM; SAHA: 1 mM in DMSO; U18: 5 µg/ml in ethanol). Control cultures run in parallel were treated with vehicle (0.1% DMSO or 0.1% ethanol or both in DMEM) for indicated times.

### Cell number and viability

Fibroblasts were cultured in 24-well plates (#833922, Sarstedt) at 15,000 cells/well and treated as indicated. Cells were detached with trypsine/EDTA (0.05%), resuspended in medium, stained with propidium iodide (2 µg/ml; Sigma-Aldrich/Merck) to label dead cells, and subjected to flow cytometry (CytoFlex, Beckman Coulter) counting all cells and propidium iodide-positive cells (excitation at 488 nm, emission 585/42 band pass filter).

### Lysate preparation and immunoblotting

Fibroblasts were cultured in 6-well plates at 150,000 cells/well and treated as indicated. Cells were lysed in homogenization buffer (sucrose 0.1M, KCl 0.05M, KH_2_PO_4_ 0.04M, EDTA 0.04M, pH 7.4, with protease inhibitor cocktail (1:1,000) and phosphatase inhibitor cocktail (1:400; Sigma-Aldrich/Merck) by sonication (VCX 130 PB, Sonics Materials) on ice for 20 sec. Then, samples were spun down at 13,000 rpm for 10 min at 4°C to remove cell debris. Protein concentrations were assessed by the Bradford method (Sigma-Aldrich/Merci) following the manufacturer’s instructions. For immunoblotting, samples were diluted with Laemmli buffer, boiled for 5 min, and subjected to SDS-PAGE (40 µg of protein/lane). Proteins were transferred to nitrocellulose membranes (Trans-Blot Turbo Transfer System; Bio-Rad Laboratories). Membranes were blocked with fat-free milk (5% in Tris-buffered saline 0.138M NaCl, 0.027M KCl, 0.025M Tris-HCl, and 0.05% Tween-20, pH 6.8) for 1h at room temperature, exposed to antibodies against NPC1 (1:1,000; #NB400-148, Novus Biologicals) or vinculin as loading control (1:40,000; #V9131, Sigma-Aldrich) overnight at 4°C followed by horseradish peroxidase-conjugated secondary IgG antibody (Bio-Rad Laboratories) for 1h at room temperature. Chemiluminescence was visualized using the ChemiDoc MP system (Bio-Rad Laboratories) and analyzed by ImageJ (National Institutes of Health, Bethesda, MD, USA).

### Endoglycosidase H Assay

Fibroblasts were cultured in 6-well plates and treated as indicated. Cells were lysed in homogenization buffer (sucrose 0.1M, KCl 0.05M, KH_2_PO_4_ 0.04M, EDTA 0.04M, pH 7.4) by sonication (VCX 130 PB, Sonics Materials) at 4°C for 20 sec, and centrifuged at 12,000 rpm for 10 min at 4°C to yield total lysate. The endoglycosidase H (EndoH) assay (V4875, Promega) was performed following the manufacturer’s instructions. The samples contained EndoH reaction buffer, water, and EndoH enzyme. As negative control, the EndoH enzyme was replaced by water. All samples were incubated at 37°C for 6 hours, and the reaction was terminated by adding Laemmli sample buffer before immunoblotting.

### Immunocytochemical staining

Fibroblasts were cultured in 96-well microplates (Black/Clear Flat Bottom Imaging Microplate; #353219, BD Falcon, Becton-Dickinson) at 3,000 cells/well and treated as indicated. Following treatment, cells were washed three times with phosphate-buffered saline (PBS), and chemically fixed with 4% paraformaldehyde (in PBS) for 15 min at room temperature. Cells were permeabilized (saponin 0.05% in PBS; #84510, Sigma-Aldrich/Merck) for 10 min, and incubated for 45 min with blocking solution (3% BSA with 1% goat serum in PBS) and then overnight at 4°C with primary antibodies (1% BSA in PBS) against NPC1 (1:2,500; NB400-148, Novus Biologicals) and LAMP2 (1:1,000; #sc-18822, Santa-Cruz). After incubation, cells were washed and reacted for 1h at room temperature with appropriate secondary antibodies (1:1,000; goat anti-rabbit secondary antibody Alexa Fluor 546; #A-10040, ThermoFisher Scientific; goat anti-mouse secondary antibody Alexa Fluor 488; #A-11001, ThermoFisher Scientific). Fluorescence was visualized and digitized using an upright microscope (Zeiss Observer 7) equipped with a light source (Zeiss Colibri), objectives (40x water, N.A. 1.2; 63x oil, N.A. 1.4; Zeiss), a module for optical sectioning by structured illumination (Zeiss ApoTome.2) and a digital camera (Hamamatsu ORCA-Flash 4.0). Densities of LAMP2- and NPC1-positive puncta were determined in manually outlined regions of interest (ROI) (1-5 per soma) using custom-written Labview (National Instruments) routines ^74^. Colocalization was estimated based on Pearson’s correlation coefficient of NPC1 and LAMP2 fluorescence intensities in individual puncta detected in ROIs.

### Cytochemical staining

Fibroblasts were cultured in 96-well microplates (black imaging plate; #353219, BD Falcon) at 3,000 cells/well and treated as indicated. Following treatment, cells were washed three times with PBS, fixed by paraformaldehyde (4% in PBS) for 15 min at room temperature and stained with filipin (50 μg/mL in PBS prepared freshly from a 250-fold ethanolic stock; #F9765, Sigma-Aldrich/Merck) for 2h at room temperature in the dark. Fluorescence images of stained cells were acquired using an inverted microscope (Axiovert 135TV; Zeiss) equipped with a metal halide lamp (10%; Lumen 200; Prior Scientific), an appropriate excitation/emission filter (XF02-2; Omega Optical Inc.), a 40x objective (oil, N.A. 1.3; Zeiss) and an air-cooled monochrome charge-coupled device camera (Sensicam, PCO Computer Optics) controlled by custom-written Labview routines (National Instruments). Densities of filipin-positive puncta and fluorescence intensities were determined in 10 to 12 images per condition and preparation from manually outlined ROIs (1-5 per cell) using custom-written Labview (National Instruments) routines^74^.

### Lysotracker staining and cytometry

Fibroblasts were cultured in 6-well plates (#833920 Sarstedt) at 150,000 cells/well and treated as indicated. Before the end of the treatment, cells were incubated with LysoTracker Red DND-99 (1 µM in culture medium; #L7528, Life technologies) for 30 minutes at 37°C, detached, spun-down at 13,000 rpm and resuspended in 300 μL of medium prior to flow cytometry. Data were acquired using a flow cytometer (Cytoflex-LX; Beckman-Coulter) and analyzed by specialized software (Kaluza, Beckman-Coulter; FloJo, Becton-Dickinson). Fold-changes in LysoTracker intensity were calculated as ratios of geometric means of stained / unstained samples as described ^75^.

### Hexosaminidase activity assay

Fibroblasts were cultured in 96-well microplates (#833924, Sarstedt) at 4,000 cells/well in DMEM without phenol red. The assay was performed similar as described (Demais et al., 2016). Briefly, following treatment, 20 µL of cell culture medium were incubated at 37°C for 3 hours with 20 µL reaction mix containing sodium citrate (10 mM; pH 4.2) and 4-methylumbelliferyl-2-acetamido-2-deoxy-b-D-glucopyranoside (2 mM; #474502, Sigma-Aldrich/Merck). The reaction was stopped by 5 volumes of glycine and Na_2_CO_3_. (0.2M). Crystal violet (#C0775, Sigma-Aldrich/Merck; 0.05% in H_2_O with 1% paraformaldehyde and 1% methanol) was added to the cell suspension to indicate cell number. The fluorescent product 4-methylumbelliferone and crystal violet were measured in triplicate on a microplate reader (Tecan Spark, Männedorf, Switzerland) using suitable filters (excitation: 365 nm; emission filters: 440 nm; absorbance: 585 nm). Calibration curves were acquired using defined amounts of the fluorescent product 4-methylumbelliferone sodium salt (#M1508, Sigma-Aldrich/Merck). Fluorescence of 4-methylumbelliferone was normalized to crystal violet intensity.

### Data analysis and visualisation

Data analysis and visualisation were accomplished with ImageJ, and with custom written routines using the open source software R (R Core Team, 2021) and selected packages (data.table, ggplot2). Unless indicated otherwise, results are displayed using bar and whisker plots representing mean and standard deviation, respectively. Statistical tests were performed as indicated. When comparing three or more experimental groups, analysis of variance (ANOVA) was carried out, followed by Tukey’s post-hoc test. Asterisks indicate statistically significant differences based on p values (*, p < 0.05; **, p < 0.01; ***, p < 0.001).

## Results

We explored BET proteins as a new therapeutic drug target for NPCD using the membrane-permeant inhibitor JQ1 and primary cultures of dermal fibroblasts from NPCD patients and a healthy donor.

### Effects of JQ1 on viability, protein levels of NPC1 and its distribution in fibroblasts

First, we tested whether the drug affected the number and viability of dermal fibroblasts. As shown in Fig. 1A, patient-derived fibroblasts carrying the I1061T variant attained lower numbers and showed a lower fraction of propidium iodide-positive (dead) cells compared to fibroblasts from the healthy donor. JQ1 inhibited cell growth compared to vehicle (DMSO) (Fig. 1B) as reported previously in dermal fibroblasts ^76^ and in immortalized cell lines ^58^. This effect occurred regardless of the genotype (Fig. 1B). The drug enhanced the percentage of dead cells at 48h of treatment in healthy fibroblasts, but more robust effects occurred in patient-derived cells after 96h of treatment (Fig. 1B).

**Figure 1.**
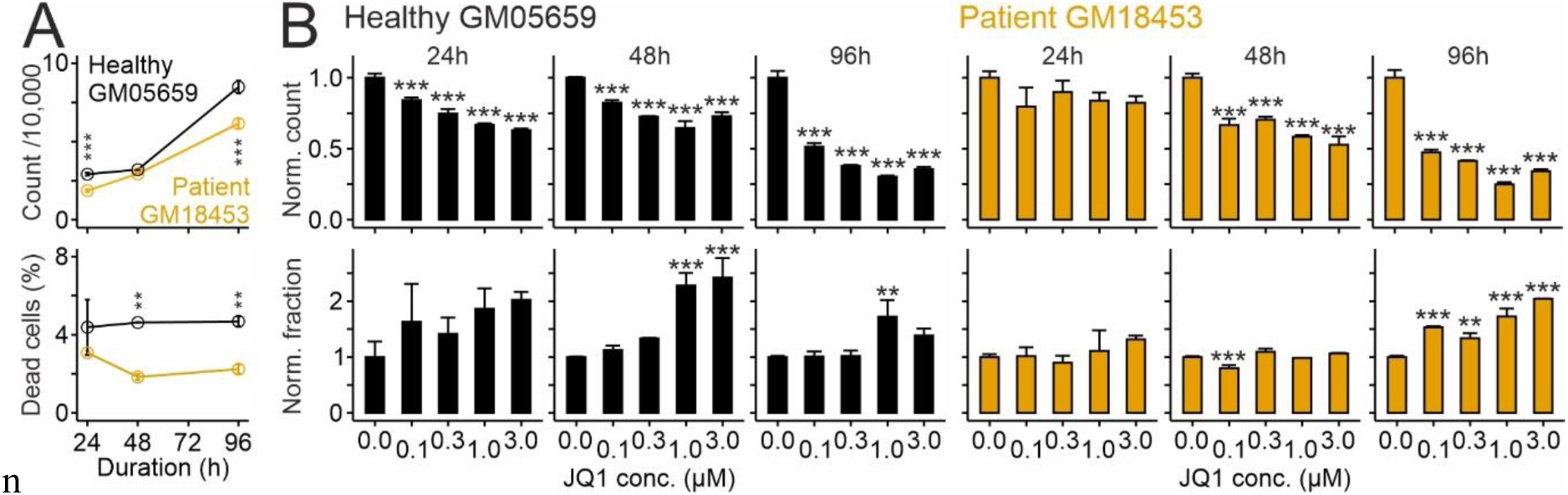
Effects of JQ1 on cell number and viability in primary cultures of human skin fibroblasts. Cell counts (top) and percentages of propidium iodide-positive cells (bottom) in primary cultures of skin fibroblasts from a healthy donor (GM056549; black) and a NPCD patient (GM018453; orange) in untreated cells (A) and after treatment with JQ1 or vehicle (DMSO) for indicated periods and concentrations (B). Values in B were normalized to values from vehicle-treated cultures run in parallel. Asterisks indicate statistically significant changes (ANOVA with Tukey’s post-hoc test; n = 3 independent preparations for each experiment).

Next, we tested how JQ1 affects NPC1 protein levels in human fibroblasts. These experiments followed up on our previous observation that the drug modifies components mediating lipid homeostasis, including NPC1 in a hepatocarcinoma cell line ^59^. Immunoblotting revealed lower levels of the I1061T variant compared to the normal version of NPC1 in fibroblasts (Fig. 2A) in line with previous studies ^63,77,78^. JQ1 enhanced protein levels of NPC1 in a concentration-dependent manner compared to vehicle-treated (JQ1 concentration zero) cultures independently from the genotype with changes reaching statistical significance after 72h of treatment (Fig. 2B). The fraction of EndoH-resistant NPC1 variant was unaffected by JQ1 (Fig. 2C) indicating that JQ1 does not modify protein glycosylation in the Golgi apparatus.

**Figure 2.**
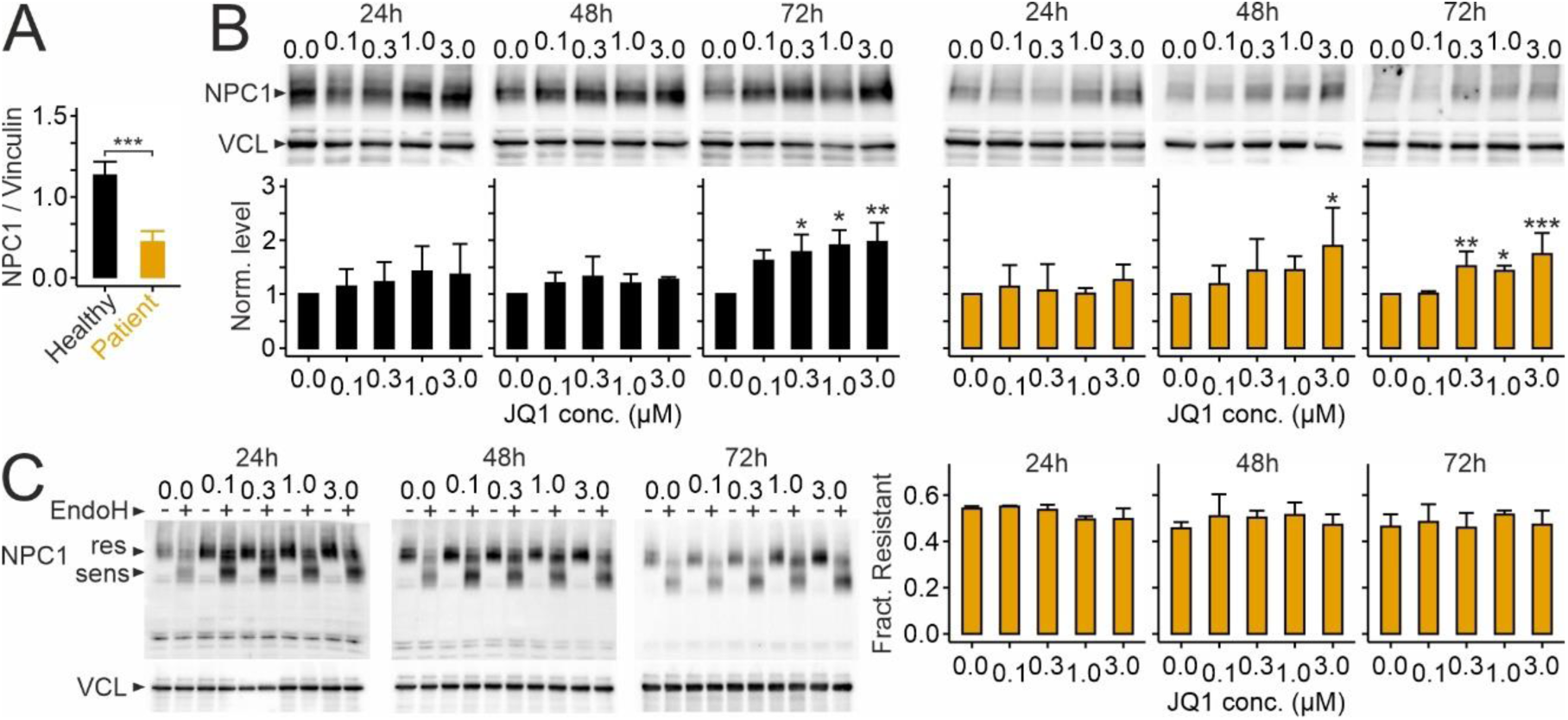
JQ1 enhances NPC1 protein levels in cultured human skin fibroblasts. A-B, Levels of NPC1 protein in primary cultures of skin fibroblasts from a healthy donor (GM05659; black) and a NPCD patient (GM018453; orange) in untreated cells (A) and after treatment with JQ1 or vehicle (DMSO) for indicated periods and concentrations (B). Values in A and B were normalized to vinculin (VCL) concentration and to vehicle-treated (JQ1 concentration zero) control cultures, respectively. Asterisks indicate statistically significant changes (A: patient versus healthy, t test, n = 4 preparations; B: treatment versus vehicle control, ANOVA with Tukey’s post-hoc test; n = 3–7 preparations). C, Fraction of EndoH-resistant protein compared to total protein in patient-derived fibroblasts after treatment with JQ1 as indicated (n = 3 preparations per treatment). Note the change in protein size following EndoH-mediated glycan removal. Images in (B) and (C) show representative immunoblots reacted with antibodies against NPC1 and VCL as loading control.

Next, we asked whether JQ1-induced NPC1 protein reaches the endosomal-lysosomal system using double immunocytochemical staining with antibodies against NPC1 and against the endosomal-lysosomal marker LAMP2 (Fig. 3A). Quantitative analyses confirmed lower levels of NPC1 and of its colocalization with LAMP2 in patient-derived fibroblasts compared to healthy donor cells (Fig. 3B). Treatment of patient-derived cells with JQ1 did not enhance the density of LAMP2 or NPC1-positive clusters (Fig. 3A, C) and only moderately affected colocalization of NPC1 and LAMP2 (Fig. 3C).

**Figure 3.**
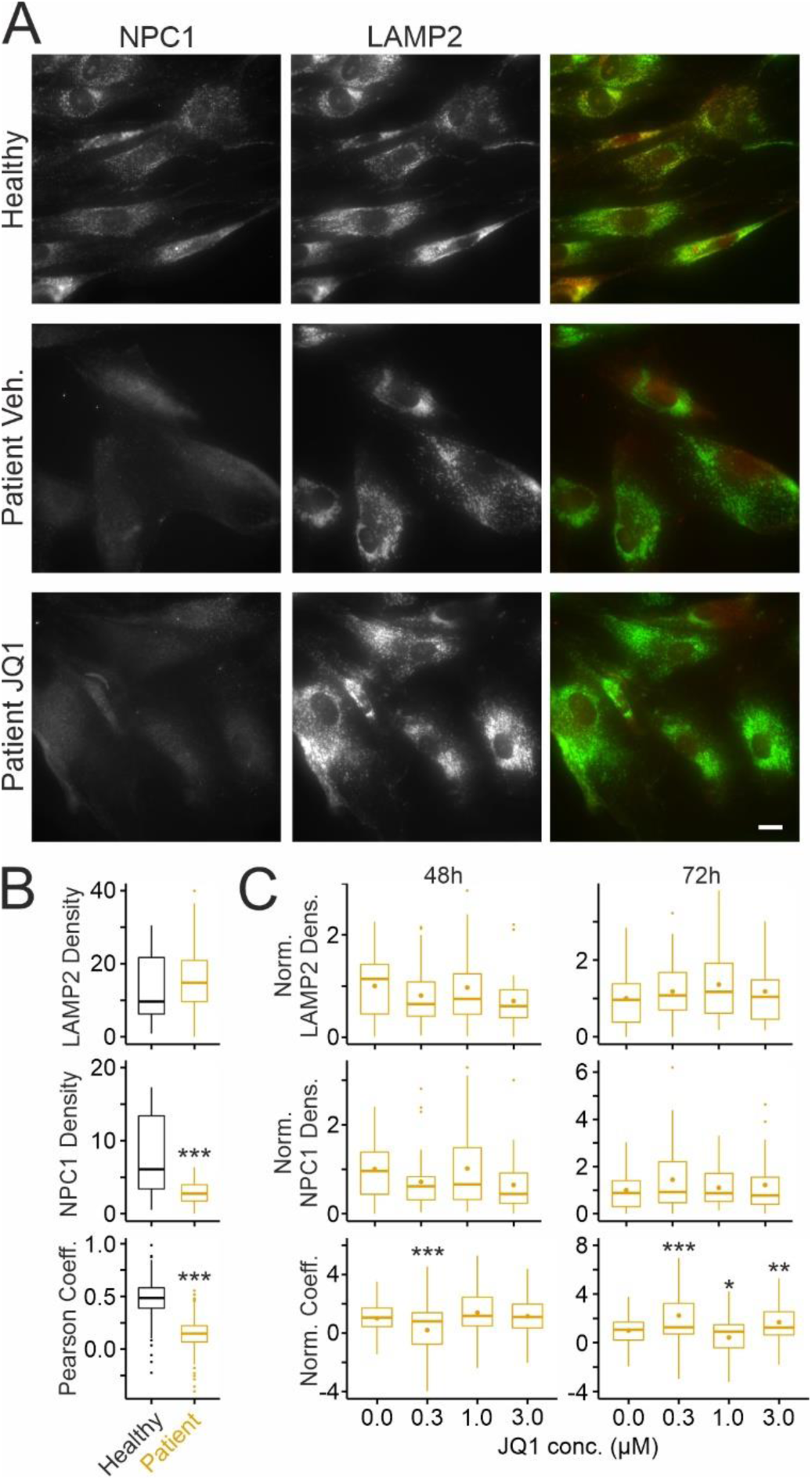
Effects of JQ1 on subcellular distribution of NPC1 in cultured human skin fibroblasts. A, Fluorescence micrographs of cultured fibroblasts from a healthy donor (top) (GM056549) and a NPCD patient (GM18453) following treatment for 72h with vehicle (DMSO) (middle) or with JQ1 (3 µM; bottom). Cells were fixed and subjected to immunocytochemical staining for NPC1 (left) and LAMP2 (middle). False-color micrographs (right) represent overlays of NPC1 (red) and LAMP2 fluorescence (green). Nuclear staining in the red channel is artificial and independent from the presence of NPC1. Scale bar: 20 µm. B-C, Boxplots showing the density of LAMP2 (top) and of NPC1-positive puncta (middle) and Pearson’s correlation coefficients of NPC1 and LAMP2 intensities (bottom) in somata of patient and healthy donor fibroblasts without treatment (B), and in patient fibroblasts following treatment with JQ1 as indicated (C). In C, respective values were normalized to the means of vehicle-treated (JQ1 concentration zero) cultures. Asterisks indicate statistically significant changes (B, top, middle: t test; n = 31/31 healthy/patient images from 3 preparations with 10-11 images per preparation and condition and 5-31 ROIs per image; bottom: n = 31/31 healthy/patient images from 3 preparations with 10-11 images per preparation and 3-29 ROIs per image; C: ANOVA with Tukey’s post-hoc test; top, middle: 48h: n = 31 to 33 images per concentration from 3 preparations with 10-12 images per treatment and 4 to 37 ROIs per image; 72h: n = 32 to 34 images per concentration from 3 preparations with 10-13 images per treatment and 4 to 37 ROIs per image; bottom: 48h: n = 31 to 33 images per concentration from 3 preparations with 10-12 images per treatment and 1 to 37 ROIs per image; 72h: n = 32 to 33 images per concentration from 3 preparations with 8-13 images per treatment and 2 to 33 ROIs per image).

### Impact of JQ1 on pathologic changes in patient-derived fibroblasts

We tested next whether JQ1 affects hallmarks of NPC1 deficiency in these cells, notably the accumulation of unesterified cholesterol and the expansion of the acidic (lysosomal) compartment, using cytochemical staining with the cholesterol-binding drug filipin ^23,79^ and with lysotracker, a pH-sensitive fluorescent dye ^75,80–82^, respectively (Fig. 4). Patient-derived fibroblasts showed a higher density of filipin-positive puncta (Fig. 4A, B), larger cell complexity and stronger lysotracker fluorescence compared to cells from the healthy donor (Fig. 4E–G). JQ1 increased the density of filipin-positive puncta after 48h of treatment (Fig. 4C) and showed no effects after 72h (Fig. 4C). However, closer scrutiny of this treatment period revealed that JQ1 induced a dual effect in a given cell population with the majority showing decreased and 30% of cells showing increased filipin densities (Fig. 4D). These divergent changes explained the lack of significance and suggested a gradual time-dependent switch of the JQ1 effect. To test this possibility, we treated cells with JQ1 for 168h. During long-term treatment, JQ1 reduced the density of filipin-positive puncta in all patient-derived fibroblasts (Figs. 4C,D) indicating time- and dose-dependent effects of the drug on cholesterol accumulation. Flow cytometry of lysotracker-labeled cells confirmed that cells carrying the disease-causing variant of NPC1 have a larger acidic compartment than cells from a healthy donor (Fig. 4E–G). Notably, treatment of patient-derived cells with JQ1 reduced this compartment in a dose-dependent manner at all time points tested (Fig. 4H).

**Figure 4.**
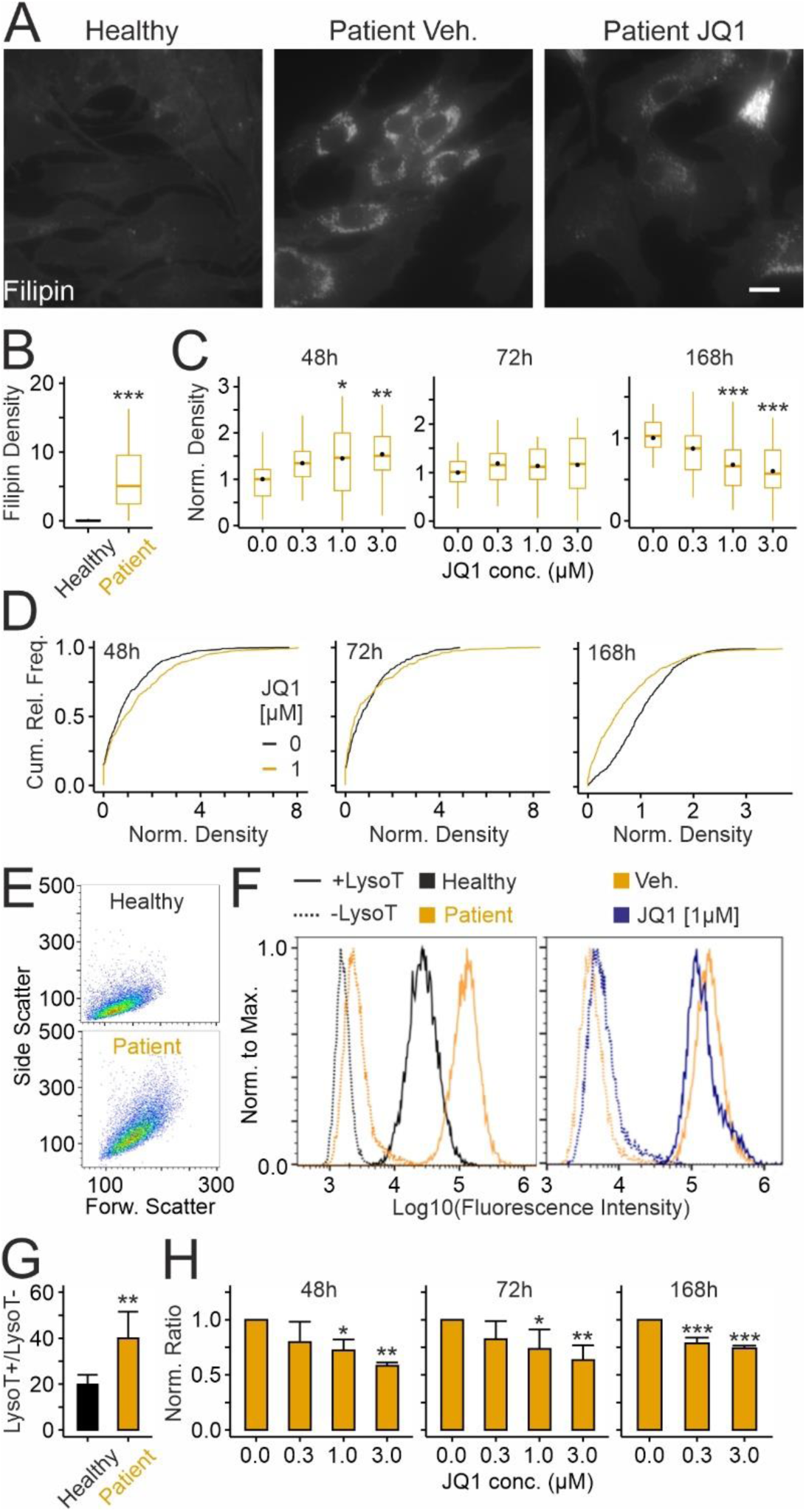
JQ1 reduces pathologic changes in lysosomes of cultured skin fibroblasts from a NPCD patient. A, Fluorescence micrographs of cultured fibroblasts from a healthy donor (GM056549) and a NPCD patient (GM18453) following treatment for 72h with JQ1 (3 µM) or with vehicle (DMSO). After chemical fixation, cells were subjected to cytochemical staining with filipin to reveal the distribution of unesterified cholesterol. Scale bar: 20 µm. B, C, Boxplots showing densities of filipin-positive puncta in fibroblasts from the healthy donor and the NPCD patient (B) and following treatment with JQ1 for indicated durations and concentrations (C). Black circles indicate mean values. Values in C were normalized to the means of vehicle-treated cultures. Asterisks indicate statistically significant changes (B: t test; n = 45/45 healthy/patient images from 4 preparations with 10-11 images per preparation and condition and 6-30 ROIs per image; C: ANOVA with Tukey’s post-hoc test; 48h: 33-34 images per concentration from 3 preparations with 10-11 images per preparation and condition and 1-24 ROIs per image; 72h: 33-34 images per concentration from 3 preparations with 10-12 images per preparation and condition and 6-27 ROIs per image; 168h: 33-34 images per concentration from 3 preparations with 10-12 images per preparation and condition and 3-32 ROIs per image). D, Cumulative relative frequency plots showing densities of filipin-positive puncta in fibroblasts from the NPCD patient (GM18453) following treatment for indicated periods with JQ1 (1 µM) or with vehicle (DMSO; 48h, vehicle: 450 ROIs / 1 µM: 462; 72h, vehicle: 527/434; 168h: 818/519; same data as shown in panel C). Note the time-dependent effects of JQ1 increasing and decreasing the densities at 48h and 168h, respectively, and the dual effect at 72h with 30 and 70% of cells showing larger and smaller densities, respectively. E, Representative plots of side and forward scatter intensities of fibroblasts from the healthy donor (top) and from the NPCD patient (bottom) obtained by cytometry. F, Histograms of fluorescence intensities normalized to maximal counts in untreated cells from the healthy donor, and the NPCD patient treated with vehicle (DMSO) or with JQ1, and unstained (dotted line) or stained with lysotracker (solid line) as indicated. G-H, Mean ratios of geometric means (lysotracker signal divided by background signal from cells without lysotracker) in untreated fibroblasts from the healthy donor and the NPCD patient (G), and in patient-derived fibroblasts following treatment with JQ1 (H) as indicated. In H, ratios were normalized to the means of DMSO-treated (JQ1 concentration zero) cultures. Asterisks indicate statistically significant changes (G: t test; n = 4 preparations; H: ANOVA with Tukey’s post-hoc test; n = 3–5 preparations).

### JQ1-induced lysosomal exocytosis

The JQ1-induced reduction of the lysotracker signal may have been due to exocytotic release of lysosomal content ^83^. To address this idea, we used two complementary assays ^74,83^. First, immunocytochemical staining of LAMP2 in non-permeabilized cells revealed the presence of LAMP2 on the cell surface following fusion of the lysosomal with the plasma membrane. Second, detection of hexosaminidase activity in the culture medium revealed the cellular release of lysosomal enzymes (Fig. 5). The assays showed that NPC1 deficiency enhances basal LAMP2 surface expression (Fig 5A, B) and extracellular hexosaminidase activity (Fig. 5D) in patient-derived fibroblasts compared to fibroblasts from a healthy donor. Treatment with JQ1 further increased both the LAMP2 surface expression (Fig. 5C) and extracellular hexosaminidase activity in a time- and concentration-dependent manner (Fig. 5E) suggesting that the drug induces release of lysosomal material.

**Figure 5.**
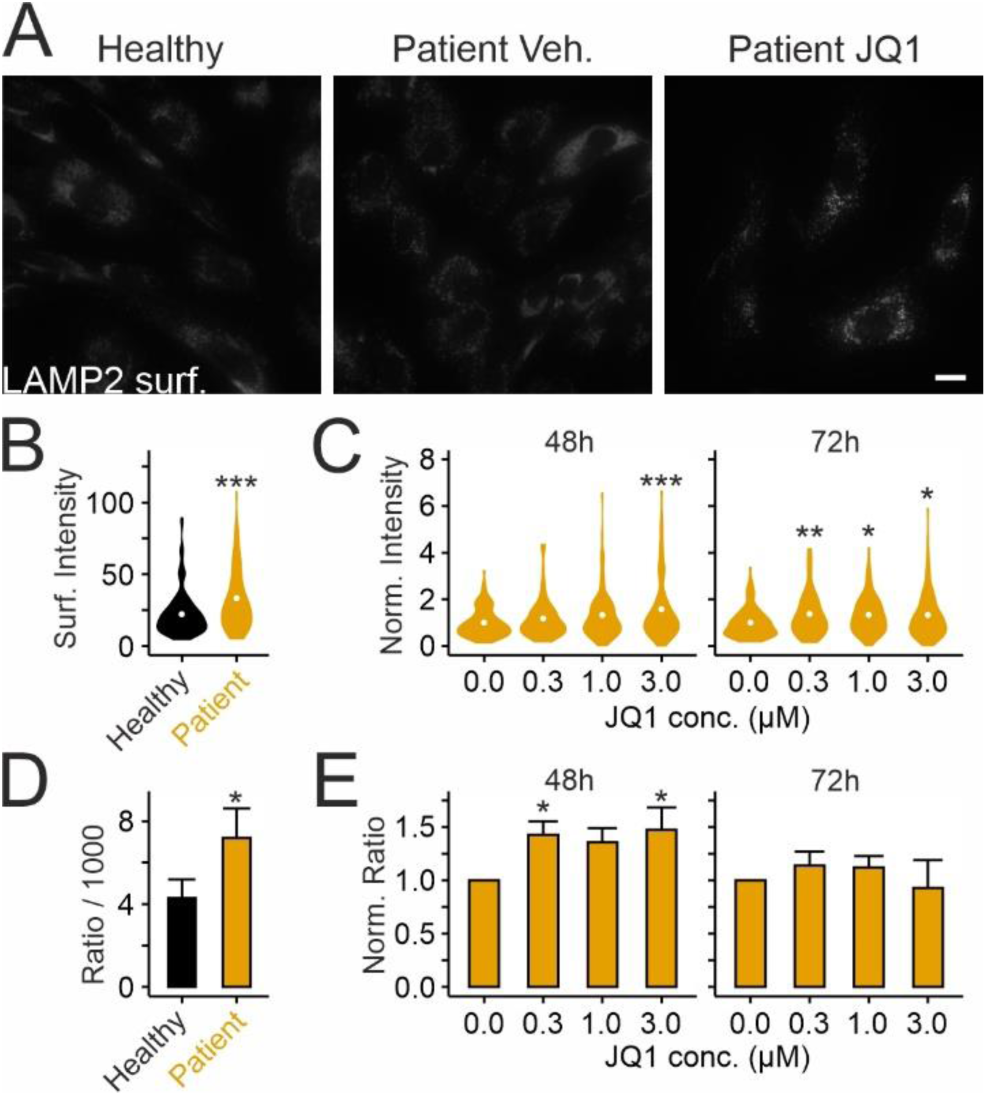
JQ1 induces extracellular release of lysosomal material in cultured skin fibroblasts from a NPCD patient. A, Fluorescence micrographs of cultured fibroblasts from a healthy donor (GM056549) and a NPCD patient (GM18453) after treatment for 72h with JQ1 or vehicle (DMSO). Following chemical fixation without permeabilization, cells were subjected to immunocytochemical staining to reveal the surface distribution of LAMP2. Scale bar: 20 µm. B-C, Violinplots showing fluorescence intensities indicating surface expression of LAMP2 on somata of fibroblasts without treatment (B), and in patient-derived fibroblasts following treatment with JQ1 as indicated (C). In C, intensities were normalized to the means of vehicle-treated (JQ1 concentration zero) cultures. White points indicate mean values. Asterisks indicate statistically significant changes (B: t test; n = 153/131 healthy/patient-derived cells from 4 preparations; C: ANOVA with Tukey’s post-hoc test; 48h: 85–116; 72h: n = 102–131 patient-derived cells per treatment from 3–4 preparations). D-E, Mean hexosaminidase activity normalized to cell numbers in untreated fibroblasts from the healthy donor and the NPCD patient (D), and in patient-derived fibroblasts following treatment with JQ1 as indicated (E). In E, values were normalized to the means of vehicle-treated (JQ1 concentration zero) cultures (D: t test; n = 3–4 preparations; E: ANOVA with Tukey’s post-hoc test; n = 3–4 preparations).

### Effects of JQ1 on fibroblasts from different patients

We next tested how JQ1 affects NPC1 levels and cholesterol accumulation in skin fibroblasts from four patients bearing different variants and polymorphisms. As shown in Fig. 6, these lines had different levels of NPC1 protein and different degrees of cholesterol accumulation under untreated conditions. Treatment with JQ1 for 72h raised the protein levels of NPC1 by two- to four-fold in a dose-dependent manner in all but one fibroblast line (GM17920). With respect to cholesterol accumulation, we observed distinct outcomes, as the effects of JQ1 differed markedly between patient-derived lines. The spectrum ranged from a strong decrease already within 72h in line GM03123 to an apparent lack of response in line GM17920. In the other lines, JQ1 reduced cholesterol accumulation after 168h of treatment (Fig. 6D). Notably, in all lines tested, JQ1 showed dual effects increasing and decreasing puncta density in subsets of fibroblasts from the same culture preparation (Fig. 6E) as shown in GM18453 (Fig. 4D). These results indicated that JQ1 robustly raises NPC1 levels in most patient-derived lines. With respect to cholesterol accumulation, subsets of cells responded to JQ1 with increased and decreased densities in a time- and dose-dependent manner with a net reduction occurring in 3 out of 4 lines.

**Figure 6.**
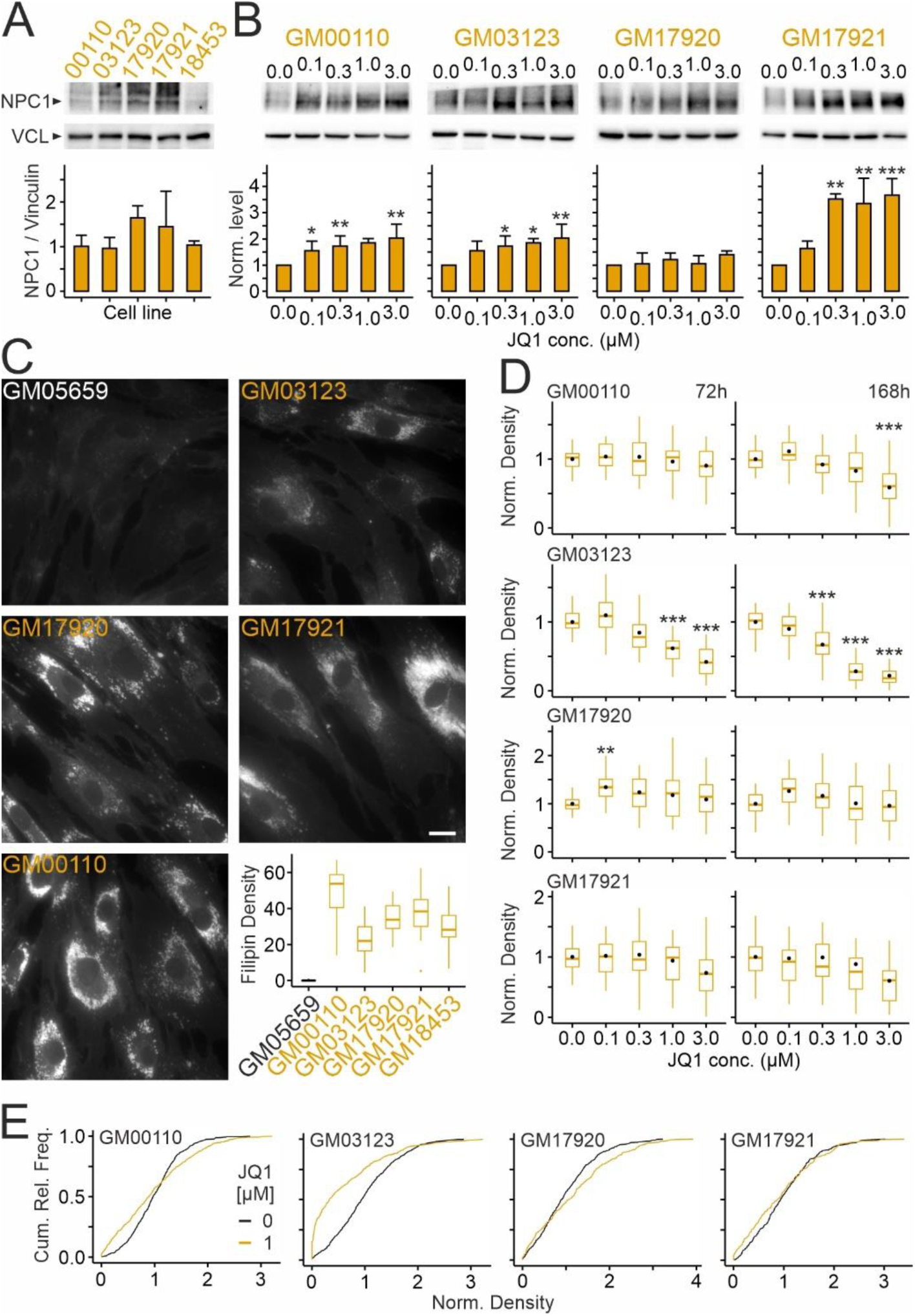
JQ1 enhances NPC1 levels and reduces cholesterol accumulation in cultured skin fibroblasts in a patient-specific manner. A,B, Levels of NPC1 protein in primary cultures of skin fibroblasts from different NPCD patients under basal levels (A) and after treatment with JQ1 or vehicle (DMSO) at indicated concentrations for 72h (B). Top, representative images of Western blots showing bands corresponding to NPC1 and vinculin. Bottom, column plots in A and B showing mean values normalized to vinculin (VCL) levels (A: n = 4 preparations) and to vehicle-treated (JQ1 concentration zero) control cultures (B: n = 3–5 preparations), respectively. Asterisks in B indicate statistically significant changes compared to vehicle control (ANOVA with Tukey’s post-hoc test). C, Fluorescence micrographs of cultured fibroblasts from the healthy donor and from several NPCD patients (indicated by codes) showing basal levels of unesterified cholesterol without treatment. Cells were subjected to chemical fixation and cytochemical staining with filipin. Scale bar: 20 µm. Boxplots showing densities of filipin-positive puncta in somata of the different fibroblast lines under untreated conditions (GM05659: n = 502 ROIs; GM00110: n = 664; GM03123: n = 568; GM17920: n = 647; GM17921: n = 487; GM18453: n = 596; 41-45 images per line; 9-12 images per preparation; 4 preparations each). D, Violin plots showing densities of filipin-positive puncta in fibroblasts from indicated patients following treatment with JQ1 at indicated durations and concentrations. Density values were normalized to the means of vehicle-treated (JQ1 concentration zero) cultures. White circles indicate mean values. Asterisks indicate statistically significant changes (ANOVA with Tukey’s post-hoc test; GM00110: 72h: n = 481–659 ROIs, 32 images per conc., 8–34 ROIs per image; 168h: 500–653 ROIs, 31–32 images per conc., 1–37 ROIs per image; GM03123: 72h: 418–673 ROIs, 31–32 images, 8–33 ROIs per image; 168h: n = 456–629, 28–32 images, 7–33 ROIs per image; GM17920: 72h: n = 329–505 ROIs, 30 images, 3–23 ROIs per image; 168h: n = 297–436 ROIs, 30 images, 4–21 ROIs per image; GM17921: 72h: 263–338 ROIs, 30 images, 4–21 ROIs per image; 168h: 249–354 ROIs, 29–30 images, 2–23 ROIs per image; 9-12 images per preparation, condition and line; n = 3 preparations per patient line). E, Cumulative relative frequency plots showing densities of filipin-positive puncta in indicated fibroblast lines following treatment for 72h with vehicle (DMSO) or JQ1 at 1 µM (subset of data shown in D). Note the dual effects of JQ1 increasing and decreasing puncta densities that occur in all patient lines.

### Effect of JQ1 on cholesterol accumulation in patient-derived fibroblasts in the presence of a NPC1 inhibitor

Our findings raised the question whether the reduction of cholesterol accumulation by JQ1 depends on NPC1 activity. To address this point, we applied JQ1 in the presence or absence of U18, an inhibitor of NPC1 ^84^, using the GM03123 line, which strongly responded to JQ1 treatment (Fig. 6D). As shown in Fig. 7, treatment with U18 enhanced the intensity of filipin fluorescence in a dose-dependent manner indicating that the cells have residual NPC1 activity. Co-treatment with JQ1 further enhanced the staining intensity at each U18 concentration tested (Fig. 7) indicating that JQ1 does not reduce cholesterol accumulation in an NPC1-independent manner.

**Figure 7.**
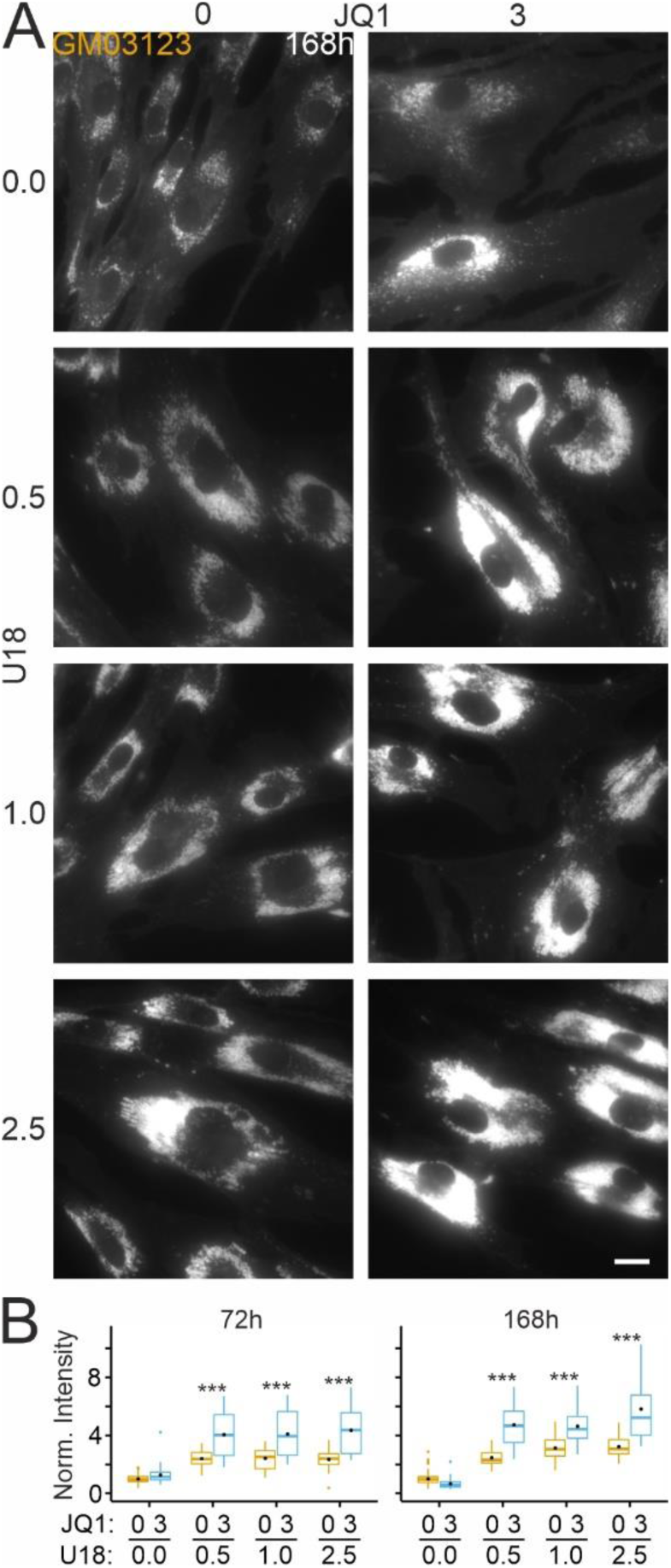
Effects of JQ1 on cholesterol accumulation in the presence of the NPC1 inhibitor U18. A, Fluorescence micrographs of cultured fibroblasts from a NPCD patient (GM03123) treated for 168h with vehicle (DMSO/ethanol) or with U18 (microgram per mL) and JQ1 (micromolar) at indicated concentrations. Following treatments, cells were subjected to chemical fixation and cytochemical staining with filipin. Scale bar: 20 µm. B, Boxplots showing fluorescence intensities of filipin staining in somata of fibroblasts treated for indicated durations with the indicated drug combinations. Intensity values were normalized to the means of vehicle-treated (JQ1 concentration zero) cultures. Black circles indicate mean values. Asterisks indicate statistically significant changes induced by JQ1 on U18-treated cells (two-way ANOVA with Tukey’s post-hoc test; 72h: n = 406–818 ROIs, 37-40 images per condition, 2–92 ROIs per image; 168h: 344–739 ROIs, 28–45 images per condition, 1–73 ROIs per image; 3 preparations).

### Effects of JQ1 on SAHA-mediated reduction of cholesterol accumulation in patient-derived fibroblasts

Previous studies revealed that pharmacologic inhibition of histone deacetylases (HDACs) reduces cholesterol accumulation in mouse neural stem cells ^85^ and human fibroblasts ^77,86,87^ carrying NPC1 variants. Therefore, we next tested whether inhibition of BET proteins, which act as histone acetylation readers, modifies these effects. In line with previous reports, the HDAC inhibitor SAHA reduced the density of filipin-positive puncta in a dose-dependent manner (Fig. 8A,B). Interestingly, similar to JQ1, also SAHA induced dual effects with fibroblasts in the same culture well showing enhanced and reduced puncta densities (Fig. 8C). Addition of JQ1 further reduced the density of puncta but the changes were not statistically significant based on p levels (Fig. 8B,C).

**Figure 8.**
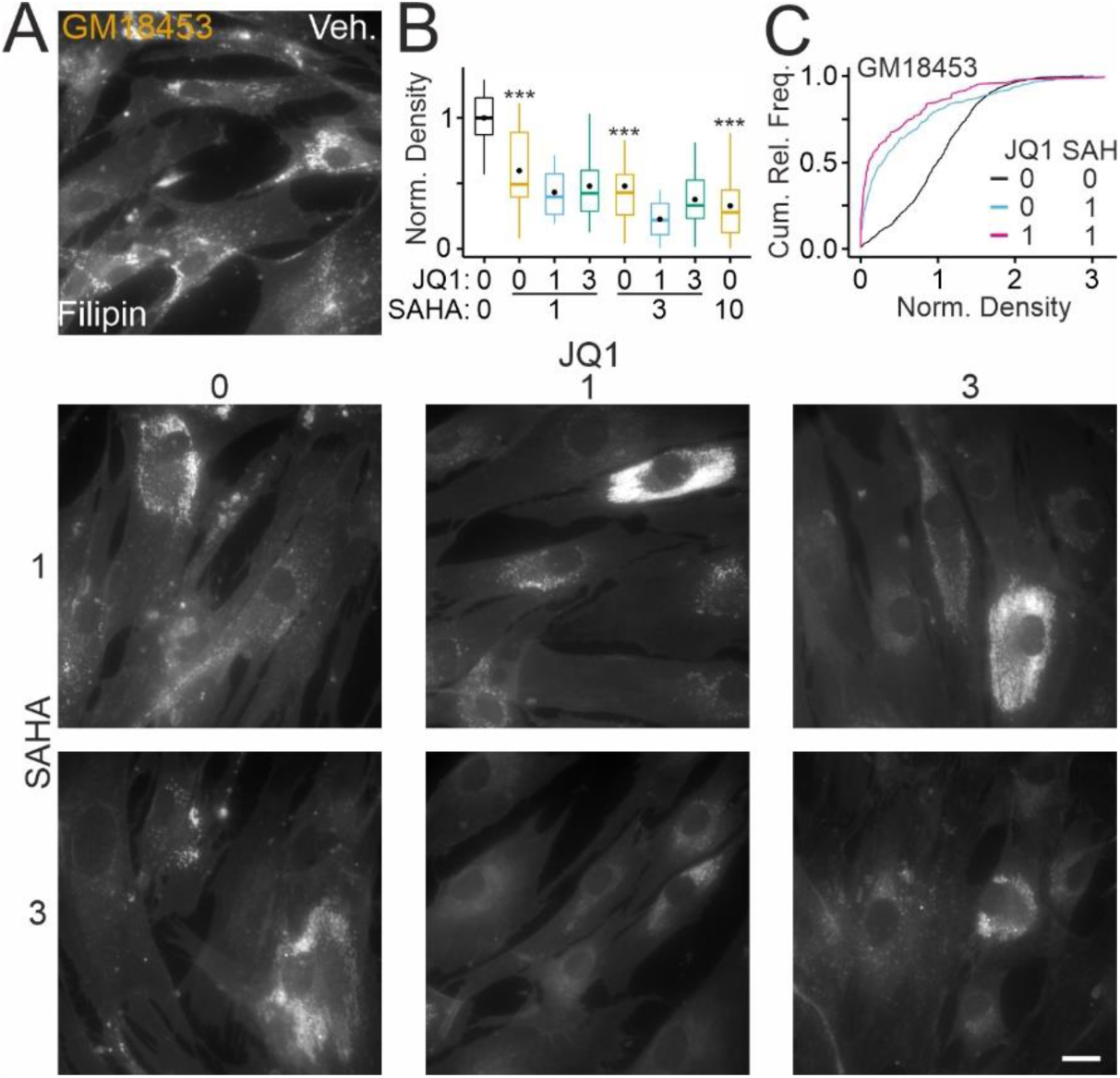
Dose-dependent effects of JQ1 on SAHA-mediated reduction of cholesterol accumulation. A, Fluorescence micrographs of cultured fibroblasts from a NPCD patient (GM18453) treated for 72h with vehicle (DMSO) only (Veh.) or with indicated combinations of SAHA and JQ1 at micromolar concentrations. Following treatments, cells were subjected to chemical fixation and cytochemical staining with filipin. Scale bar: 20 µm. B, Boxplots showing densities of filipin-positive puncta in somata of fibroblasts treated with the indicated drug combinations. Co-treatment with JQ1 and 10 µM SAHA was toxic to cells precluding further analyses. Density values were normalized to the means of vehicle-treated (JQ1 concentration zero) cultures. Black circles indicate mean values. Asterisks indicate statistically significant changes induced by SAHA compared to vehicle-treated cells (two-way ANOVA with Tukey’s post-hoc test; n = 368–605 ROIs, 29–40 images per conc., 6–29 ROIs per image. C, Cumulative relative frequency plots showing densities of filipin-positive puncta in the patient-derived fibroblast line following indicated treatments for 72h. Note the dual effect of SAHA similar as observed for JQ1 (Figs. 4D, 6E), and additive effects of SAHA and JQ1. Subsets of data shown in panel B.

## Discussion

Here, we report previously unknown effects of BET protein inhibition in a cell culture model of NPCD that encourage further exploration of this drug candidate. We found that a prototypical BET protein inhibitor enhanced the cellular level of NPC1 protein, diminished lysosomal expansion and cholesterol accumulation, and induced the release of lysosomal components in a time- and dose-dependent manner. The effects of JQ1 on cholesterol levels depended on NPC1 activity, the enhancement of protein levels occurred in most of the patient lines tested, but the extent of cholesterol reduction varied in a line-dependent manner. JQ1 reinforced the reduction of cholesterol accumulation induced by the HDAC inhibitor SAHA.

Our finding that BET protein inhibition enhances protein levels of NPC1 is in line with previous reports that the promoter region of *Npc1* is associated with acetylated histones ^88^ and that de-acetylation increases gene expression ^86^. More generally, BET proteins have been shown to control lysosome- and autophagy-related genes in various cell types and disease conditions ^53,89–96^. Other means to enhance cellular NPC1 protein levels have been explored *in vitro* including inhibition of histone deacetylases ^65,68,77,97–99^, enhanced chaperone activity ^63,72,78,100–106^, and reduced protein degradation ^63,64,100,102^. These manipulations enhanced the presence of NPC1 variants in the endosomal-lysosomal system ^63,64,68,72,78,99–101,104,105^, and they reduced the intracellular accumulation of unesterified cholesterol ^63–65,68,77,78,86,87,98–101,104,105^. Our observation that JQ1 bolstered the effect of SAHA on cholesterol accumulation suggests that treatment with drugs affecting distinct epigenetic pathways may be beneficial. There is evidence for synergistic effects of BET protein and HDAC inhibitors in transcription regulation ^107^ and tumor therapy ^108,109^. Dosing of each drug probably requires adjustment to avoid unwanted effects ^110,111^.

Our observation that the effect of JQ1 on cholesterol accumulation varies in fibroblasts from different patients is in line with line-specific effects of HDAC inhibitors ^68,77,99^. The variable outcome probably depends on the specific residual activity of the variant ^63,68,99^. This is further supported by our finding that JQ1 failed to reduce cholesterol accumulation after pharmacologic inhibition of NPC1 by U18. Evidently, other genetic or epigenetic modifiers ^112–114^ may further impact the outcome of JQ1 treatment. Notably, we observed dual effects of JQ1 on cholesterol accumulation in primary fibroblasts with their net balance changing from an early increase to a net decrease after prolonged treatment. The mixed response observed at 72h with some cells showing an increase and others a decrease may explain the seeming lack of effect of JQ1 on cholesterol accumulation in a recent fibroblast-based high-throughput drug screen for NPCD ^115^. The time-dependent reversal of JQ1 effects on cholesterol accumulation may be caused by sequential activation of distinct processes or subtype-specific responses ^116^. The initial increase of cholesterol accumulation may be due to enhanced levels of lysosomal components as reported previously ^89^ and the sparse integration of NPC1 in the endosomal-lysosomal system. During this period, JQ1 also reduced the lysosomal volume as indicated by lysotracker staining. This effect may have been caused by immediate release of lysosomal content to the extracellular space as indicated by the enhanced presence of LAMP2 on the cell surface and increased hexosaminidase activity in the medium. Lysosomal exocytosis reduces the extent of cholesterol accumulation due to NPC1 dysfunction as suggested by previous studies on cell lines ^117–121^, patient-derived fibroblasts ^117,122–124^, and retinal neurons ^74,125^. The net decrease of cholesterol accumulation after prolonged JQ1 treatment may be caused by additional processes that are affected by BET protein inhibition ^126^, including a reduction of cholesterol biosynthesis ^59,127^ and of intracellular lipid levels ^59^, and an increase of apolipoprotein a ^128^, which has been explored as therapeutic agent for NPCD ^129^. In general, BET protein inhibition affects many transcriptional programs through interactions with acetylated lysines on histones including a site showing epigenetic marks in NPCD ^130^.

Taken together, our results reveal that BET proteins regulate NPC1 levels and thereby impact cholesterol homeostasis depending on specific protein variants and the genetic background.

## Author contributions

Conceptualization: AS, FWP, MS, VP; Data curation: AB, AS, FWP, MP, SL, VP; Formal Analysis: AB, AS, CT, FWP, MP, SC, VP; Funding acquisition: FWP, MS, VP; Investigation: AB, AS, CT, MP, SC, SL; Methodology: AB, AS, CT, FWP, LS, MP, MS, SC, SL, VP; Project administration: AB, CT, FWP, MP, VP. Resources: AS, FWP, VP; Software: AS, FWP; Supervision: FWP, VP; Validation: AB, FWP, MP, VP; Visualization: AS, FWP; Writing – original draft: FWP; Writing – review & editing: AB, AS, FWP, MP, MS, SC, SL, VP.

## Abbreviations

ANOVA: analysis of variance
BET: bromodomain and extraterminal domain
DMSO: dimethylsulfoxide
EndoH: endoglycosidase H
HDAC: histone deacetylase
NPCD: Niemann-Pick type C disease
SAHA: suberoylanilide hydroxamic acid

## Ethics/integrity statements

### Data availability

All data are contained within the manuscript. Data will be shared upon request to FWP or VP.

